# Universal constraints on selection strength in lineage trees

**DOI:** 10.1101/2020.12.04.412106

**Authors:** Arthur Genthon, David Lacoste

## Abstract

We obtain general inequalities constraining the difference between the average of an arbitrary function of a phenotypic trait, which includes the fitness landscape of the trait itself, in the presence or in the absence of natural selection. These inequalities imply bounds on the strength of selection, which can be measured from the statistics of traits or divisions along lineages. The upper bound is related to recent generalizations of linear response relations in Stochastic Thermodynamics, and is reminiscent of the fundamental theorem of Natural selection of R. Fisher and of its generalization by Price. The lower bound follows from recent improvements on Jensen inequality and is typically less tight than the upper bound. We illustrate our results using numerical simulations of growing cell colonies and with experimental data of time-lapse microscopy experiments of bacteria cell colonies.

## Introduction

Quantifying the strength of selection in populations is an essential step in any description of evolution. With the development of single cell measurements, a large amount of data on cell lineages is becoming available both at the genotypic and phenotypic level. By analyzing the statistics of cell divisions in population trees, one can measure selection more accurately than using classical population growth rate measurements [1]. Similarly, by tracking phenotypes on cell lineages, one can obtain statistically reliable estimations of the fitness landscape of a given trait and of the selection strength of that trait [2]. All these methods contribute to bridging the gap between single-cell experiments at the population level and molecular mechanisms [3].

An alternate method to infer selection in evolution focuses on dynamical trajectories of frequency distributions [4, 5]. In these works, Mustonen et al. introduced the notion of fitness flux to characterize the adaptation of a population by taking inspiration from Stochastic Thermodynamics. In fact, ideas from Stochastic Thermodynamics can be applied directly at the level of individual cell trajectories [6]. By following this kind of approach, we have derived general constraints on dynamical quantities characterizing the cell cycle such as the average number of divisions or the mean generation time [7, 8]. These constraints are universal because they hold independently of the specific cell dynamic model and they are indeed verified in experimental data. Other examples of universality in the context of evolution includes the identification of universal families of distributions of selected values and the use of methods from extreme value statistics [9, 10].

Here, we derive universal constraints for the average value of a trait, and for its selection strength by exploiting a set of recent results known under the name of Thermodynamic Uncertainty Relations (TUR). These relations take the form of inequalities, which generalize fluctuation-response relations far from equilibrium [11], and which capture important trade-offs for thermodynamic and non-thermodynamic systems [12] as recently reviewed in [13]. Although our results are framed in the context of cell population in lineage trees, they apply more broadly to general stochastic processes defined on any branched tree.

### Framework

A colony of cells can be represented as a branched tree, whose branches are called lineages and whose nodes correspond to cell divisions. We assume that each cell in the population divides after a stochastic time into exactly *m* daughter cells. In order to extract relevant statistics from such a tree, one needs to sample the lineages following a weighting scheme. Two main weighting methods have been introduced in [2, 3, 14], namely the backward (or retrospective) and the forward (or chronological) samplings. The backward sampling of lineages assigns a uniform weight *N* (*t*)^*−*1^ to each of the *N* (*t*) lineages, leading to an over-representation of cells coming from sub-populations that divided more than average. To compensate this bias, the forward sampling takes into account the number of divisions *K* along a lineage and assigns to the lineage a weight 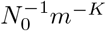, where *N*_0_ is the size of the initial population. It cancels the effect of natural selection acting on a colony, and allows to extract from tree-structured data the statistics that would be obtained in single-lineage experiments, like in mother-machine configuration [15].

The statistical bias is captured by a relation between the two probability distributions for the number of divisions, similar to fluctuation relations in Stochastic Thermodynamics [7, 8]

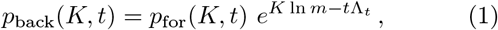

where Λ_*t*_ = ln(*N* (*t*)*/N*_0_)*/t* is the population growth rate. When the population reaches a steady-state exponential growth, Λ_*t*_ tends to a constant growth rate Λ.

By the same logic, one defines the fitness landscape of the value *s* of a general phenotypic trait 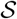 by *h*_*t*_(*s*) [2] using the following relation between snapshot probability distributions at time *t*

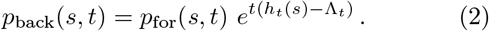

This relation quantifies the selection bias acting on individuals carrying the trait value *s* in a population. Note that *h*_*t*_(*s*) is related to but distinct from the growth rate of the subpopulation carrying the trait value *s* as explained in Suppl. Mat. Importantly, *h*_*t*_(*s*) could be called fitness seascape instead of fitness landscape since it is typically time-dependent [5]. Keeping that in mind, we stick with the original terminology below.

Two constraints on the population growth rate follow from eq. (2) by considering the positivity of the Kullback-Leibler (KL) divergences 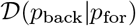 and 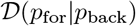 of the two probability distributions [2]

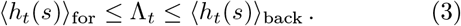

To go beyond these inequalities, the exact value of Λ_*t*_ may be obtained by integrating eq. (2) over *s*, which leads to

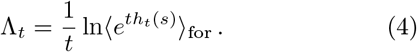

Note that eq. (4) provides an estimator for the population growth rate Λ_*t*_, which requires only mother machine data [8]. This is useful, although in practice the convergence of this estimator is a challenging issue [16], which is one motivation to look for inequalities as done below.

### Fluctuation-response inequality for populations of cells

Let us introduce an arbitrary function of the value *s* of a general trait 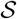 as *g*_*t*_(*s*). Then, we define the ratio of the backward to forward probabilities as

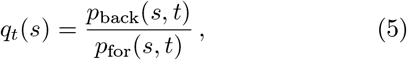

and consider the product 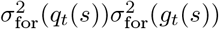, with 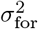 the variance with respect to the forward distribution. From now on, we omit the dependence of functions *q*_*t*_ and *g*_*t*_ in *s* in order to make the notations lighter. We then use the Cauchy-Schwarz inequality to obtain

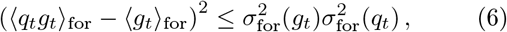

where we used 〈*q*_*t*_〉_for_ = 1 due to the normalization of *p*_back_. Finally, using the relation 〈*q*_*t*_*g*〉_for_ = 〈*g*_*t*_〉_back_ and taking the square root of our expression, we obtain

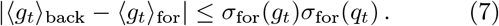

Note that that the l.h.s. of eq. (7) involves averages with respect to the two probability distributions unlike what happens in the standard TUR where only one such average is present. The reason is that in Stochastic Thermodynamics, the two relevant probability distributions correspond to a forward and a time-reversed dynamics, and the quantity which replaces *g*_*t*_(*s*) is a current, which changes sign under time reversal symmetry. Here there is no such symmetry present, hence the two averages are not the opposite of one another.

The inequality can be understood as an out-of-equilibrium generalization of fluctuation-dissipation equalities [11], which involves a comparison between a reference unperturbed dynamics (the forward sampling) and a perturbed dynamics (the backward sampling), with selection as the perturbation. In our case, the difference between the unperturbed and the perturbed averages of the function *g*_*t*_(*s*) is bounded by the unperturbed fluctuations of this function, measured by *σ*_for_(*g*_*t*_), times *σ*_for_(*q*_*t*_) which is a ‘pseudo-distance’ between the two probability distributions similar to a KL divergence. Indeed, since 〈*q*_*t*_〉_for_ = 1, the variance of *q*_*t*_ is given by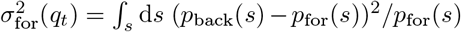, and thus the larger *σ*_for_(*q*_*t*_), the further away *p*_back_(*s, t*) and *p*_for_(*s, t*) are from each other.

While this interpretation is general, the ‘pseudo-distance’ term *σ*_for_(*q*_*t*_) can be linked to measurable quantities for cell colonies. Indeed, using eqs. (2) and (4), we obtain

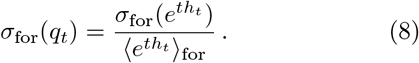

Thus, the distance term quantifies the relative fluctuation of the quantity 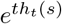, which itself represents the ratio of the expected number of lineages ending with trait value *s*, rescaled by the number *N*_0_ of initial cells, to the forward probability of this trait value value (See Suppl. Mat.).

### Linking fluctuations of fitness landscape to the strength of selection

An important application of the above result is when the arbitrary function *g*_*t*_(*s*) is the fitness landscape *h*_*t*_(*s*) itself. Then, the change in the mean fitness in the ensemble with and without selection represents the strength of selection 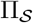 acting on trait 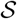 [2]

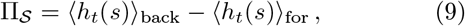

which is always positive due to eq. (3). For this choice of function *g*_*t*_(*s*), eq. (7) reads

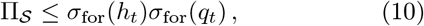

With words, eq. (10) means that the strength of the selection of any trait is bounded by the fluctuations of fitness landscape in the dynamics without selection times the distance between the perturbed and unperturbed distributions for this trait. Some more comments are in order: first, in eq. (7) and eq. (10), we have used the forward probability distribution as reference, but similar bounds can be obtained in terms of backward standard deviations with the replacement of *q*_*t*_(*s*) by the opposite ratio *r*_*t*_(*s*) = 1*/q*_*t*_(*s*), as detailed in Suppl. Mat. Second, the strength of selection defined here should not be confused with the coefficient of selection, usually defined as the relative difference in fitness associated with two values of a phenotypic trait [4]. Third, the strength of selection is a function of time, since fitness landscapes are time-dependent. Only if a steady state is reached in the long time limit, then *h*_*t*_(*s*) tends to a constant equal to the steady state population growth rate Λ, and the strength of selection tends to 0, as expected since selection no longer shifts trait frequencies.

### From the inequality to a linear response equality

Let us now investigate precisely the conditions for which the previous inequalities become saturated. It is straight-forward to show that when the forward and backward statistics are equal, inequalities eqs. (7) and (10) are saturated. Indeed, the l.h.s terms are 0 and the r.h.s terms are null because they contain the standard deviation of the constant quantity *q*_*t*_(*s*) = 1. We now study the case where the two probability distributions approach each other, which corresponds to the limit *tσ*(*h*_*t*_) → 0 (see Suppl. Mat.), referred to as the small variability limit.

In this limit, we obtain the l.h.s of eq. (7) as

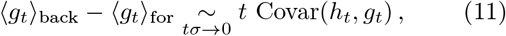

and the l.h.s of eq. (10) when the function *g*_*t*_ is the fitness landscape itself

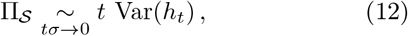

where the variance or the covariance can be equivalently taken over the forward or backward sampling. When computing the r.h.s of eqs. (7) and (10), we obtain that eq. (10) is saturated in this limit whereas eq. (7) is not.

The limit can also be written *t ≪ σ*(*h*_*t*_)^*−*1^ where *σ*(*h*_*t*_)^*−*1^ appears as a characteristic time of the system. Note that this characteristic time is itself time-dependent, since it is the inverse of the amplitude of variation of the fitness seascape, which is by definition a time-evolving landscape. In practice, this limit can be reached either for short times or in the case of a strong control mechanism on the divisions, leading the lineages to stay synchronized even after a finite time. It is also possible to regard this limit as a regime of weak selection [17], since the strength of selection is small precisely because of eq. (10).

### Gaussian approximation

In the particular case of Gaussian distributions, we show that eq. (12) stays valid beyond the small variability limit. Setting a Gaussian forward distribution with mean (*h*_*t*_)_for_ and variance 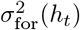 for the fitness landscape *h*_*t*_(*s*), one obtains with no expansion needed, for a bijective function *h*_*t*_(*s*) (see Suppl. Mat.)

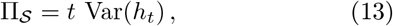

where the variance can be indifferently taken over the forward or backward sampling. Note that we recover here a result known from [2], in a more direct way and with restricted assumptions, since in that reference the authors assumed that the joint distribution of *h*_*t*_(*s*) and *K* was a bivariate Gaussian distribution.

### Connection to Fisher’s fundamental theorem

Fisher’s fundamental theorem of natural selection states that the time derivative of the mean fitness of a population is equal to the genetic variance of the fitness across the population [18]. We note that eqs. (12) and (13) are similar to Fisher’s relation, but with some differences: for instance, in the l.h.s of eqs. (12) and (13) the time derivative of mean fitness is replaced by the strength of selection for a given phenotypic trait.

One of the main limitations of Fisher’s theorem lies in the implicit assumption that natural selection is the only possible phenomenon leading to a change in the gene frequencies [19]. This assumption neglects many important phenomena such as mutations and recombination events [17], random drift due to finite population size, and specific features of seascapes [5]. In contrast, our result does not suffer from any of these limitations. Indeed, eq. (12) and eq. (13) hold for a finite population in seascapes; and the origin of the variability of cell divisions does not matter in our model, it can be partly genetic due to mutations or recombination events, and partly non-genetic. The strength of selection is built to isolate the effect of selection from other potential sources of variability, and has the advantage to be a trait-dependent measure.

Note that eq. (11) can be directly compared with the co-variance term representing the separate effect of selection in the Price equation [19].

### A lower bound for the strength of selection

To complement the upper bound on the strength of selection given by eq. (10), we now derive a lower bound which improves the trivial bound, which is zero due to eq. (3). Such a lower bound presents an interest to quantify the minimal effect of selection on a particular trait. The positivity of the strength of selection comes from the positivity of the KL divergence, itself relying on Jensen’s inequality. Therefore by improving upon Jensen’s inequality [20], one obtains better lower bounds. Let us detail how these bounds may be used to constrain the strength of selection.

We define the convex functions *φ*_for_(*x*) = *e*^*tx*^, *φ*_back_(*x*) = *e*^−*tx*^ and the function

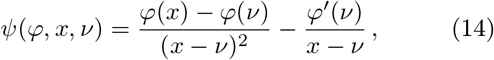

where *φ*′ stands for the derivative of *φ*. The sharpened version of Jensen’s inequality then gives (see Suppl. Mat.)

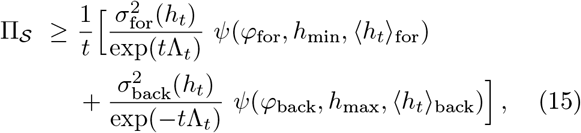

which shows that the lower bound depends on the first two moments of the forward and backward fitness landscape distributions and also on the minimal (resp. maximal) values of these distributions denoted *h*_min_ (resp. *h*_max_). When the fitness landscape is a monotonic function of the value of the trait, which is the case for cell age and size [8], or for the number of divisions, these extrema values are given by the extremal trait values.

The two terms in the r.h.s. of eq. (15) are separately positive, therefore one has a non-trivial lower bound even if only forward (resp. backward) statistics is available.

### Tests of the linear response relations

We now illustrate the various bounds for growing cell populations, using either simulations or time-lapse video-microscopy experimental data [21].

First, we tested eq. (7) for the number of divisions 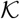, and for the linear function *g*_*t*_(*K*) = *K*, so that the inequality bounds 〈*K*〉_back_ − 〈*K*〉_for_. We simulated lineage trees starting from one cell, for a particular agent-based model in which cells are described by their sizes. Cell sizes continuously increase at constant rate between divisions, and cells divide after a stochastic time only depending on their sizes. Each simulation of such a tree yields a single point on the scatter plot fig. 1, which shows the ratio of *σ*_for_(*K*)*σ*_for_(*q*_*t*_) to 〈*K*〉_back_ − 〈*K*〉_for_ versus the population growth rate Λ_*t*_. Two sets of points are presented, which only differ in the final time of the simulation. As expected from eq. (10), all points in both sets are above 1. When the duration of the simulation is small (*t* = 3), the final population is small, around *N* ~ 20, therefore for a given tree the lineages do not have time to differentiate significantly and the variability in the number of divisions among the lineages is small. In that case, simulations points are approaching the horizontal dashed line at *y* = 1 corresponding to the saturation of the inequality. The final population *N* fluctuates significantly from one simulation to the next, because the simulation time is short and all simulations start with a single cell with random initial size. As a result, the dispersion of values of Λ_*t*_ is large.

**Figure 1:**
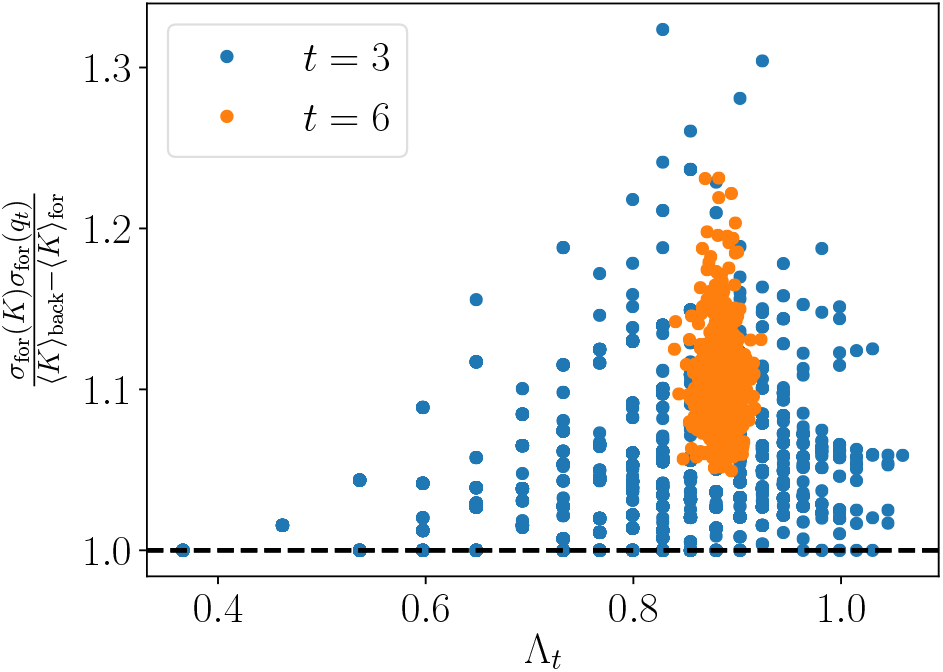
Points of *σ*_for_(*K*)*σ*_for_(*q*_*t*_)*/*(〈*K*〉_back_ − 〈*K*〉_for_) against Λ_*t*_ for many tree simulations using a size-controlled model. Each dot corresponds to a single tree, the two sets of data have the same parameters except for the final times of the simulation, which are *t* = 3 (blue) and *t* = 6 (orange). The black horizontal dashed line at *y* = 1 represents the point where the inequality of eq. (10) is saturated.

Now, when doubling the duration of the simulation, the cloud of scattered points is considerably reduced in both directions. The horizontal dispersion reduces because as *t* increases, the state of the system at the final time becomes less and less affected by the initial condition. On the vertical axis, there is a gap between the lower part of the scatter plot and the horizontal line at *y* = 1 due to the increase of heterogeneity in the number of divisions in the lineages with the simulation time.

We now test our results on real data, extracted from [21], for the following traits: the number of divisions 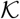, the size 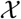 and the age 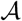, which are easily accessible and for which we predicted theoretical fitness landscapes [8]. The first step is to obtain the fitness landscapes of these traits, which are shown for three particular experimental conditions in Suppl. Mat. Remarkably, we recover for several experiments plateaus which we had predicted theoretically for the fitness landscape of cell size, and we also obtain good agreement with the general shape of the curve when the plateaus are blurred by noise.

Then, we test the upper and lower bounds on the strength of selection of cell size using this data. We show on fig. 2 the upper bound 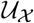 given by eq. (10) and the lower bound 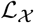 given by eq. (15), normalized by the strength of selection 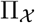. The *x*-axis numbers the colonies which have grown in different nutrient medium [21]. As expected, all the sets of points are respectively above and below the horizontal dashed line at *y* = 1. Experiments for which the normalized upper bound approaches 1 indicate that cell cycles are almost synchronized and thus that there is small variability in terms of number of divisions among the lineages.

**Figure 2:**
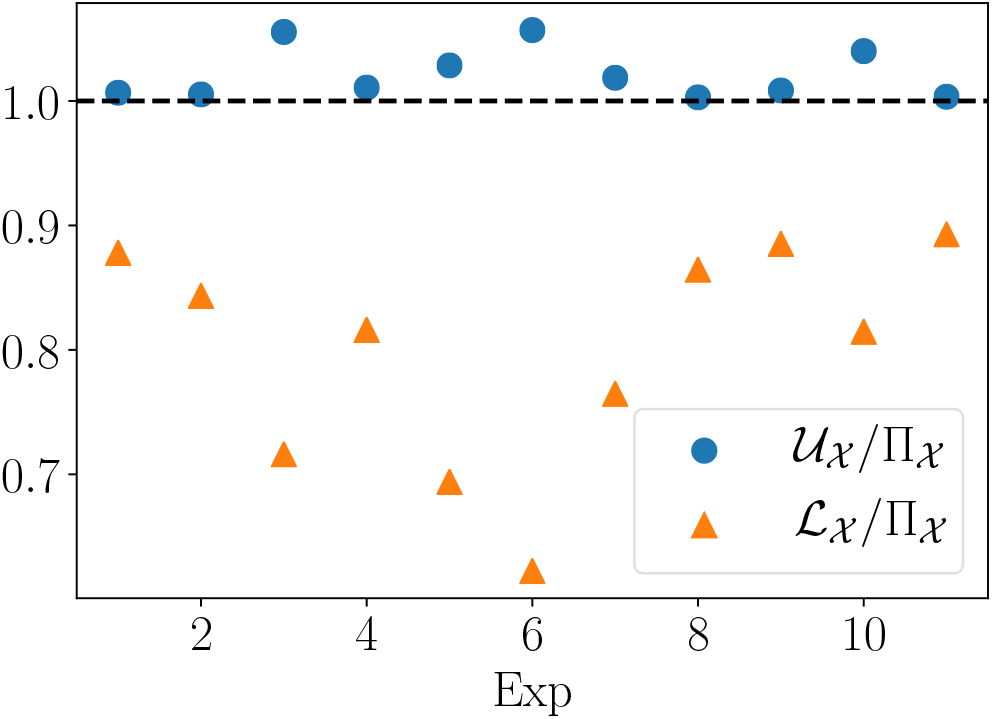
Upper bound 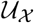 (blue dots) and lower bound 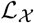 (orange triangles) for the strength of selection acting on size 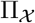, normalized by the latter. The *x*-axis represents the 11 colonies in different growth conditions from [21], in no particular order.

Note that Nozoe et al. also proved [2] that the strength of selection for the division bounds the strength of selection acting on any trait: 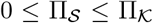. This bound, is typically not as tight as eq. (10) (see Suppl. Mat. for comparison). To improve upon it, one can use 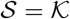 in eq. (10) to obtain a bound for 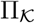 itself.

## Conclusion

The general idea of comparing the response of a system in the presence of a perturbation to its fluctuations in the absence of the perturbation lies at the heart of the Fluctuation-Dissipation theorem, which has a long history of physics, with some applications to evolution [1, 22]. Remarkably, the present framework with forward (unperturbed) and backward (perturbed) dynamics can be conveniently applied to population dynamics without having to perform additional experiments, since both probabilities can be calculated with the same lineage tree. Our main result is a set of inequalities for the average of an arbitrary function of a trait or for its fitness landscape, valid beyond the Gaussian assumption, and which constrain the strength of selection in population dynamics even in the presence of time-dependent selection pressures. For applications, we focused on phenotypic traits, like cell size or age, and in these cases, we found our upper bound on the strength of selection to be tight. In future work, it would be interesting to extend this framework to genotypic traits instead of phenotypic ones [17] and possibly exploit recent methods of lineage tracking [23]. This could open new perspectives to address a number of important problems like antibiotic resistance, the differentiation of stem cells or virus evolution.

The search for universal principles in evolution is an active field of research [22, 24]. An important step in this endeavor was made by Fisher, who boldly compared his theorem to the second law of thermodynamics. While the theorem turned out not to be as general as expected, Fisher had nevertheless the correct intuition about its importance for evolutionary biology, and he was also correct in expecting that such a general principle should be related to thermodynamics.

DL acknowledges insightful discussions with L. Peliti, O. Rivoire and J. Unterberger and support from Agence Nationale de la Recherche (ANR-10-IDEX-0001-02, IRIS OCAV and (ANR-11-LABX-0038, ANR-10-IDEX-0001-02).

## Supplementary Materials

### PHENOTYPIC FITNESS LANDSCAPE AND GROWTH RATE ASSOCIATED WITH THE VALUE *s* OF A TRAIT 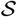

The fitness landscape *h*_*t*_(*s*) of trait 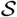 that we defined in the main text should not be confused with the fitness associated with the value *s* of the trait, which can be identified with the growth rate of the subpopulation carrying that trait, but they are linked by a simple relation.

On the one hand, in line with the definition of the population growth rate, we define the growth rate of the subpopulation carrying the value *s* of the trait 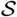 as

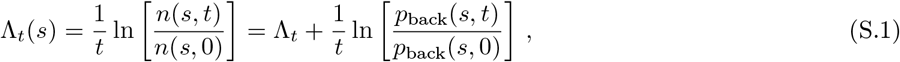

where *n*(*s, t*) = *p*_back_(*s, t*)*N*(*t*) is the number of cells with trait value *s* at time *t*. Of course, defining this quantity only makes sense between two moments *t* = 0 and *t* for which the value *s* is present in the population, otherwise the expression in undefined.

On the other hand, the definition of the fitness landscape eq. (2) is

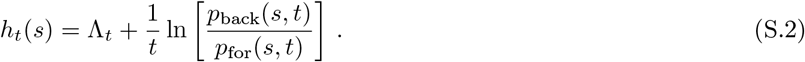

By comparing the two relations above, we can link the growth rate associated with a trait value *s* to the fitness landscape of this trait value *s*:

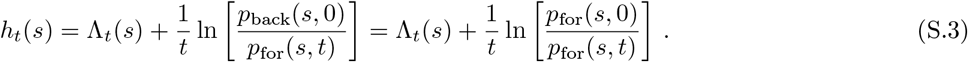

The last equality follows from the fact that, at *t* = 0, the cells have not divided yet and so the forward and backward samplings of the population are identical.

Finally, we see that the three quantities Λ_*t*_, Λ_*t*_(*s*) and *h*_*t*_(*s*) are different but intimately linked by the three relations above. In the long time limit, for which equilibrium distributions do not depend on time anymore, they all become equal to the steady-state population growth rate Λ

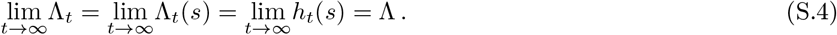

Moreover, as we already noted, eq. (S.2) indicates that the sign of *h*_*t*_(*s*) − Λ_*t*_ informs us on the comparison between the backward and forward probabilities of trait value *s* in the population at time *t*, or in other words, if the trait value *s* is over-represented in the population as compared to a situation without selection. Thus, *h*_*t*_(*s*) − Λ_*t*_ quantifies the effect of selection alone.

Following the intuitive understanding we have of the growth rate associated with a trait value *s*, eq. (S.1) means that the sign of Λ_*t*_(*s*) − Λ_*t*_ informs us on the comparison between the backward statistics of the trait value *s* at time *t* and at time 0. A trait value is favored by the population dynamics, which includes all phenomena leading to changes in trait value frequencies, if its growth rate is larger than the population growth rate, which corresponds to an increase of the frequency of that trait value in the population as time grows.

The last relation, eq. (S.3) provides a new insight: the sign of *h*_*t*_(*s*) − Λ_*t*_(*s*) is also linked to the comparison between the forward probability of trait value *s* at time 0 and time *t*. The forward statistics is constructed to balance the effect of selection occurring in tree-structured data. However, it is affected by all the other sources of variability, as for example mutations. Therefore, the sign of *h*_*t*_(*s*) − Λ_*t*_(*s*) indicates the evolution of the frequency of the value *s* of trait 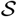 as time grows, due to every phenomenon but selection. If at division, the value *s* of a trait 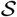 is allowed to change from the mother cell to the daughter cell, which we call mutation, and if there are statistically more mutations going from other values *s*′ to *s* than going from *s* to other values *s*′, then we say that value *s* of trait 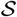 is favored by mutations, and its frequency will increase in the forward statistics, leading to *h*_*t*_(*s*) *<* Λ_*t*_(*s*).

Of course, in the same way that eq. (S.2) can be written in the form of a fluctuation theorem eq. (2), it is also the case for eqs. (S.1) and (S.3), which offers two new fluctuation theorems. A notable difference is that, unlike eq. (2), these new fluctuation relations both link two probability distributions that may not have the same support.

### ‘PSEUDO-DISTANCE’ BETWEEN PERTURBED AND UNPERTURBED DYNAMICS IN TERMS OF MEASURABLE QUANTITIES

We show in this section how the ‘pseudo-distance’ *σ*_for_(*q*_*t*_) is linked to measurable quantities in cell colonies, for a general trait 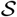, and for the particular case of age models. We showed in the main text (eq. (8)) that *σ*_for_(*q*_*t*_) is the coefficient of variation in the forward statistics of the quantity

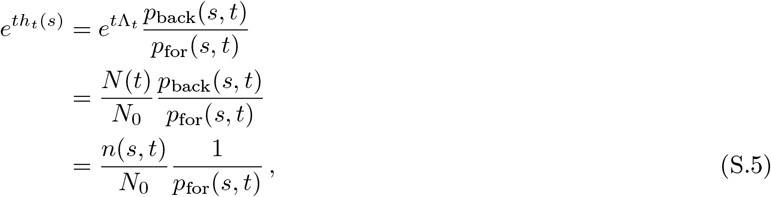

where *n*(*s, t*) = *N* (*t*)*p*_back_(*s, t*) is the number of cells with the value *s* of trait 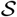 at time *t*.

In the case where 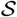 is the number of divisions, the fitness landscape is called the lineage fitness and is given by *h*_*t*_(*K*) = *K* ln *m/t*. Therefore, *σ*_for_(*q*_*t*_) = *σ*_for_(*m*^*K*^)*/ 〈m*^*K*^〉_for_ is the relative fluctuation of quantity *m*^*K*^, representing the expected number of lineages that underwent *K* divisions, normalized by the initial population *N*_0_, divided by the forward probability of *K*.

For age models, where the division is only controlled by the age of the cell, we know a fluctuation relation linking the forward and backward distributions of generation times *τ*, defined as the time between two consecutive divisions on the same lineage [25]

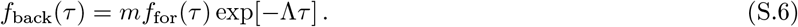

This relation can be understood as a version of the fluctuation relation on the number of divisions eq. (1) at the scale of the cell cycle. However, let us precise two differences: first, unlike eq. (1), eq. (S.6) is only true in the long time limit, when the population is growing at a constant steady state growth rate Λ; and second, the distributions *f*_back_ and *f*_for_ are not snapshot distributions at time *t*, but distributions of generation times computed along the weighted lineages.

We define *q*(*τ*) = *f*_back_(*τ*)*/f*_for_(*τ*) in the same way, and following the same steps for a general function *g*(*τ*) we derive

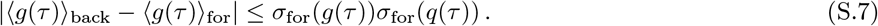

From eq. (S.6), we express the pseudo-distance term as

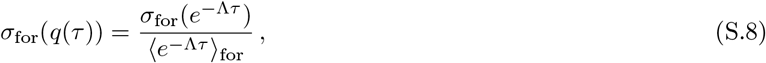

which is the relative fluctuation in the forward sampling of the quantity *e*^*−*Λ*τ*^ = *f*_back_(*τ*)*/mf*_for_(*τ*). We know from [26] that the backward distribution is also the generation time distribution of the direct ancestor cells. Therefore, *e*^*−*Λ*τ*^ represents the ratio of the probability for the ancestor cell to divide at age *τ* to the expected number of daughter cells born from that division that divide at age *τ*.

### LINEAR-RESPONSE INEQUALITY IN TERMS OF THE FLUCTUATIONS IN THE PERTURBED DYNAMICS

In this section, we show that the upper bound for the difference between the average values of an arbitrary function *g* of a phenotypic trait 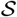, in unperturbed (forward) and perturbed (backward) dynamics, can be expressed in terms of fluctuations in the perturbed dynamics. For that purpose, we define the ratio of the forward to backward probabilities of the value *s* of trait 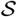:

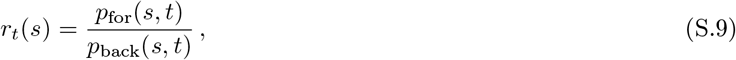

which is the inverse of the function *q*_*t*_(*s*) defined in the main text. We then follow the same steps, using the Cauchy-Schwarz inequality on the product 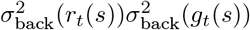, which reads

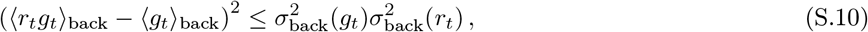

where we used 〈*r*_*t*_〉_back_ = 1 coming from the normalization of *p*_for_. Finally, using the relation 〈*r*_*t*_*g*_*t*_〉_back_ = 〈*g*_*t*_〉_for_ and taking the square root of our expression, we obtain

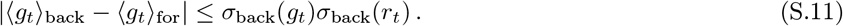

In this context, the term *σ*_back_(*r*_*t*_) = (∫_*s*_ d*s* (*p*_back_(*s*) − *p*_for_(*s*))^2^*/p*_back_(*s*))^1/2^ is still interpreted as a ‘pseudo-distance’ between the probability distributions *p*_for_(*s, t*) and *p*_back_(*s, t*), and is still connected to measurable quantities for cell colonies. Indeed, eq. (2) can be written as

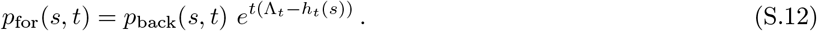

Integrating eq. (S.12) over *s* leads to

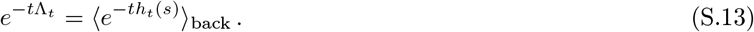

Finally, by combining eqs. (S.9), (S.12) and (S.13), we obtain

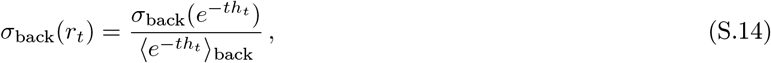

Thus, the distance term is the coefficient of variation in the perturbed dynamics of the quantity 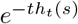. This quantity represents the ratio of the forward probability distribution *p*_for_(*s, t*) of trait value *s* to the expected number of lineages *n*(*s, t*) = *N*(*t*)*p*_back_(*s, t*) ending with this trait value, rescaled by the size *N*_0_ of the initial population:

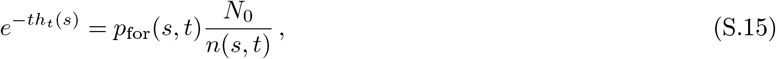

which follows directly from eq. (S.12), when using the definition of the population growth rate Λ_*t*_ = ln(*N*(*t*)*/N*_0_)*/t*.

### SMALL VARIABILITY LIMIT

In this section, we study the two sides of the fluctuation-response inequality on an arbitrary function *g*_*t*_ of a phenotypic trait 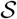 (eq. (7)) in the limit where the forward and backward distributions approach each other, and show that they are mathematically equivalent in this limit in the case where the function *g*_*t*_ is the fitness landscape *h*_*t*_. The difference between the two distributions is captured by the ‘distance’ term *σ*(*q*_*t*_(*s*)), or equivalently *σ*(ln *q*_*t*_(*s*)), where the standard deviations can be taken either in the backward or forward statistics. Using eqs. (2) and (5), this term can be reshaped as *σ*(ln(*q*_*t*_(*s*)) = *σ*(ln(*p*_back_(*s, t*)*/p*_for_(*s, t*))) = *tσ*(*h*_*t*_(*s*)). From now on, we refer to the limit where the forward and backward distributions are close to each other as the small variability limit, defined by *tσ*(*h*_*t*_(*s*)) → 0.

We first use this limit in the forward statistics: *tσ*_for_(*h*_*t*_(*s*)) → 0. Starting from the fluctuation relation on trait 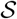 (eq. (2)), we multiply both sides by *g*_*t*_(*s*) and integrate over *s*, leading to the expression of the backward average of function *g*_*t*_ as the forward average of a biased version of the same function:

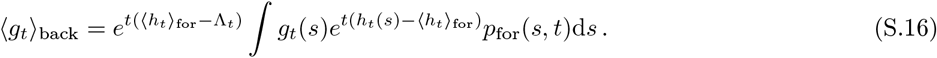

In order to expand the exponential, we assume that for any *s*, *t*(*h*_*t*_(*s*) − 〈*h*_*t*_〉_for_) is small, which corresponds to *tσ*_for_(*h*_*t*_) small because *σ*_for_(*h*_*t*_) is the characteristic distance to the mean. Therefore,

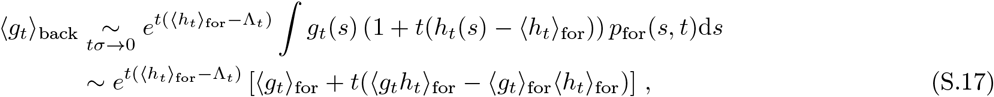

where 〈*g*_*t*_*h*_*t*_〉_for_ − 〈*g*_*t*_〉_for_〈*h*_*t*_〉_for_ = Covar_for_(*h*_*t*_, *g*_*t*_) is the covariance of *h*_*t*_ and *g*_*t*_ with respect to the forward probability. The term in the bracket is a first-order correction to 〈*g*_*t*_〉_for_ in *tσ*_for_(*h*_*t*_). Now we need to compute the prefactor exp[−*t*Λ_*t*_], starting with eq. (4) and using the same expansion

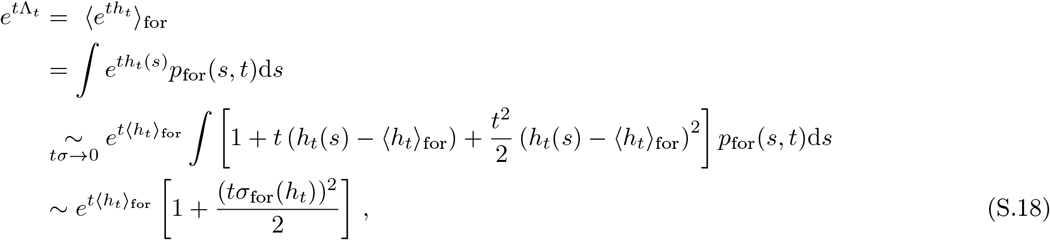

which is a second-order correction to exp[*t*〈*h*_*t*_〉_for_] in *tσ*_for_(*h*_*t*_).

Combining eqs. (S.17) and (S.18) we find that

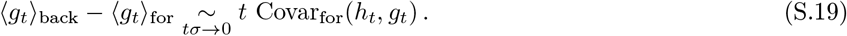

For the r.h.s. of the inequality eq. (7) of the main text, using the expression of *σ*_for_(*q*_*t*_) as the forward coefficient of variation of the quantity exp[*th*_*t*_(*s*)] (eq. (8)), it is straight-forward to show from the same expansion as the one used to obtain eq. (S.18), that

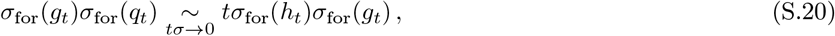

In the particular case where *g*_*t*_(*s*) is the fitness landscape *h*_*t*_(*s*), then eq. (S.19) reads

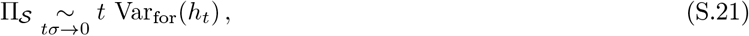

and thus the inequality eq. (10) is saturated in this limit. In contrast to that, note that because of eq. (S.19), the inequality eq. (7) does not get saturated in this limit.

As mentioned at the beginning of this section, we can then re-derive the results above from the backward point of view, using an first-order expansion in *tσ*_back_(*h*_*t*_(*s*)). In this case,

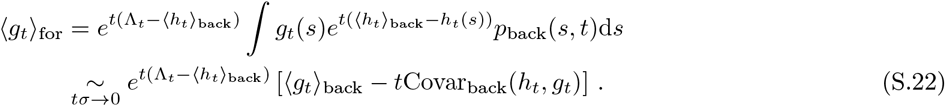

The prefactor is computed with the same expansion starting from eq. (S.13):

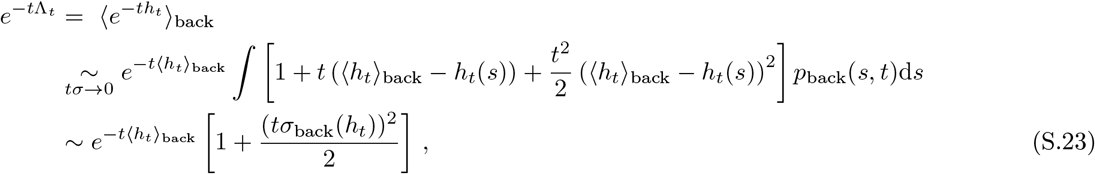

which is a second order correction in *tσ*_back_(*h*_*t*_). Combining eqs. (S.22) and (S.23), we find

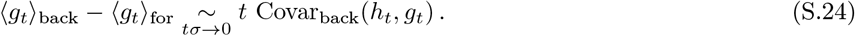

When comparing eqs. (S.19) and (S.24), we conclude that the covariance can be taken equivalently in the forward or backward statistics.

### GAUSSIAN CASE

We assume that *h*_*t*_(*s*) can be accounted for by a continuous probability distribution, even though the trait 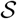 may be discrete, as it in the case for the number of divisions. We set a Gaussian forward distribution with mean 〈*h*_*t*_〉_for_ and variance *σ*_for_(*h*)^2^ for the fitness landscape *h*_*t*_(*s*), then exp(*th*_*t*_(*s*)) follows a log-normal distribution of mean

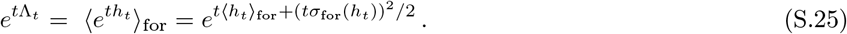

This relation shows that for a given forward average fitness landscape, the growth rate is positively affected by the variability between the lineages.

The backward average of the fitness landscape is given by the forward average of a biased fitness landscape:

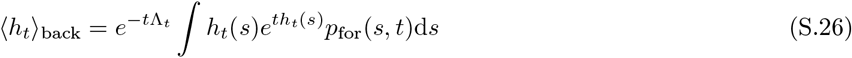

We make the hypothesis that the fitness landscape is a bijective function of the trait value and use the conservation of the probability: *p*_for_(*s, t*)d*s* = *p*_for_(*h*)d*h*, leading to a solvable Gaussian integral

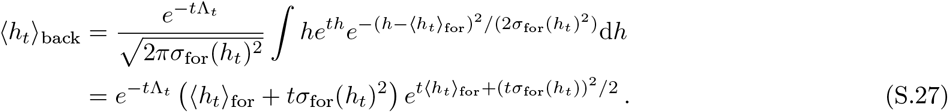

Finally, combining eqs. (S.25) and (S.27), we obtain

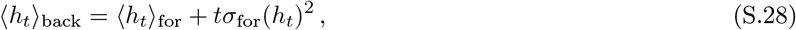

and thus

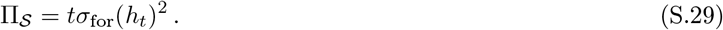

Moreover, combining eqs. (S.25) and (S.29) we deduce that 〈*h*_*t*_〉_for_ and 〈*h*_*t*_〉_back_ are not only respectively smaller and greater than Λ_*t*_, as implied by eq. (3), but they are actually symmetrical around this value 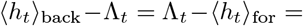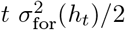. In other words, in this particular case, the KL divergence is symmetrical: 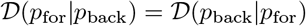.

In the case where *h*_*t*_(*s*) follows a Gaussian distribution in the forward statistics, it also follows a Gaussian distribution in the backward statistics because the bias of the fluctuation relation is exponential. We showed in eq. (S.28) the relation between the average values taken with respect to both statistics, and we aim now to show that the standard deviations of the two distributions are equal. Let us start with eq. (S.13), which links the population growth rate to the backward statistics of the fitness landscape. Since *h*_*t*_(*s*) follows a Gaussian distribution of mean 〈*h*_*t*_〉_back_ and standard deviation *σ*_back_(*h*_*t*_) in the backward statistics, then 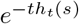 follows a log-normal distribution of mean

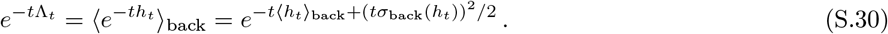

We now take the inverse of this formula and use eq. (S.28) to replace the backward average:

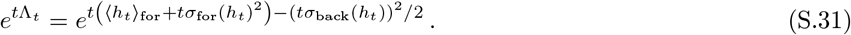

By comparing eqs. (S.25) and (S.31), it follows that *σ*_back_(*h*_*t*_) = *σ*_for_(*h*_*t*_). Finally, the standard deviation in eq. (S.29) can be taken indifferently with respect to both statistics and we omit the index to write the general version of eq. (S.29):

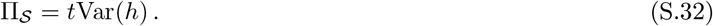

### COMPARISON WITH NOZOE’S UPPER BOUND

Nozoe et al. proved that the strength of selection acting on any trait 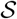 is bounded by the strength of selection acting on the number of divisions 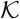 [2]: 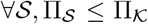. We obtained in this paper a trait-dependent bound, which highlights the role of fluctuations of fitness landscape of that particular trait for the strength of selection, but which is often tighter than 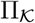. We show on fig. 3 the ratio of 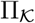 to the upper bound 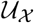 for the size (left plot) and 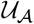 for the age (right plot). All points are indeed above 1.

**Figure 3:**
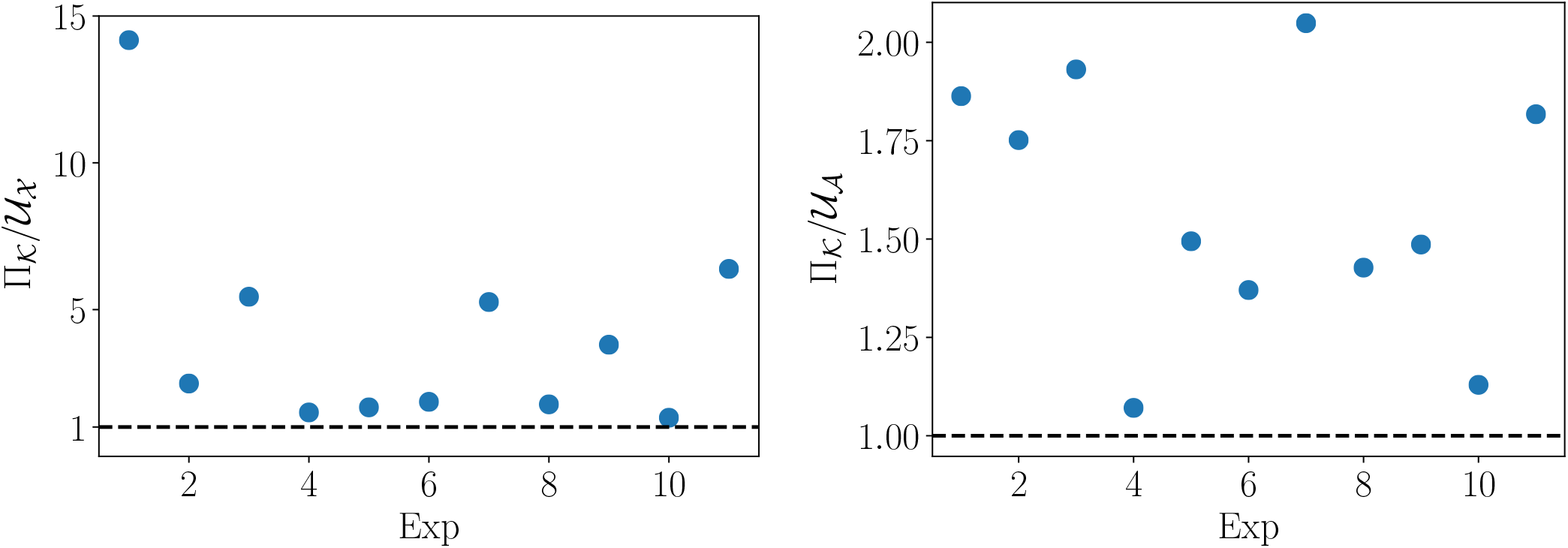
Ratio of the general bound 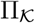 to our upper bound 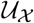 for size (left plot) and 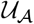 for age (right plot) for the 11 experiments from [21], in no particular order. All points are above the black horizontal dashed line at *y* = 1, which indicates that our upper bound is always smaller and thus better than 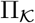.

### OBTAINING THE LOWER BOUND FOR THE STRENGTH OF SELECTION

The positivity of KL divergences 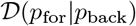 and 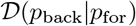, gives respectively Λ_*t*_ − 〈*h*_*t*_〉_for_ ≥ 0 and 〈*h*_*t*_〉_back_ − Λ_*t*_ ≥ 0. Then we proceed in two steps, applying Liao’s result [20] to both parts. First, seeking a positive lower bound for Λ_*t*_ − 〈*h*_*t*_〉_for_, the sharpened version of Jensen’s inequality reads

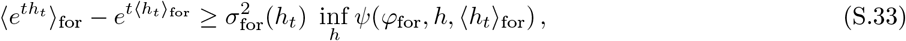

where functions *ψ*, *φ*_for_ and *φ*_back_ have been defined in the main text. The r.h.s involves the minimum of the function *ψ* when varying *h* on its support at time *t*. We then divide this expression by exp(*t*Λ_*t*_)

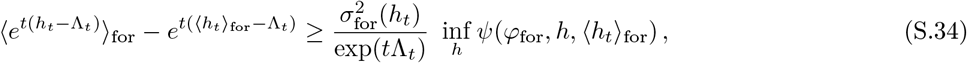

The first term in the l.h.s is given by the integration on *s* of the fluctuation relation for trait 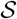 eq. (2), and its value is 1 because of the normalization of the probability distribution *p*_back_. Finally the enhanced bound reads

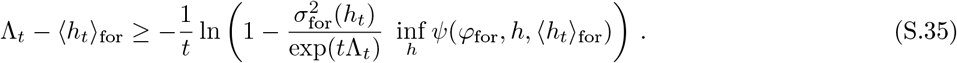

Similarly, we find

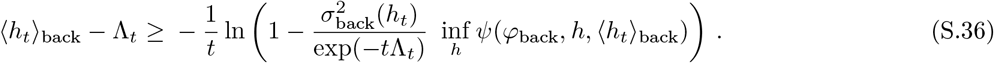

Liao et al. proved [20] that when *φ*′(*x*) is a convex (resp. concave) function, then *ψ*(*φ, x, ν*) is an increasing (resp. decreasing) function of *x*, and thus the infimum of function *ψ*(*φ, x, ν*) on *x* is reached for *x* = *x*_min_ (resp. *x* = *x*_max_).

Because of the convexity of 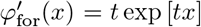, the minimum of *ψ* is reached when evaluating *ψ* at the minimum value *h*_min_ of the fitness landscape *h*_*t*_(*s*). At finite time, the support of *h*_*t*_(*s*) is finite and so is its minimum value. Similarly, because of the concavity of 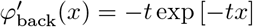, the minimum of *ψ* is reached when evaluating *ψ* at the maximum value *h*_max_. Finally, we use the relation − ln(1 − *x*) ≥ *x* valid for any real number *x* and we combine the two inequalities to obtain

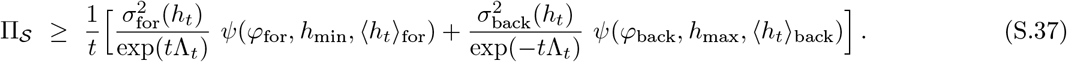

Note that the r.h.s. of eqs. (S.35) and (S.36) are both positive numbers due to the convexity and concavity of 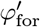 for and 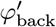 respectively. As a result, their sum is also positive and therefore the r.h.s. of eq. (S.37) does represent an improvement with respect to the trivial bound which would be 0.

Liao et al. also proposed another lower bound, looser but simpler than the one involving the function *ψ*. Indeed, one can replace inf_*h*_ ψ(*φ*_for_*, h,* 〈*h*_*t*_〉_for_) by 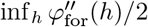 in eq. (S.33). Moreover 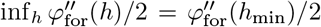 since 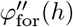 is an increasing function of *h*. The same goes for the other inequality, and combining the two leads to

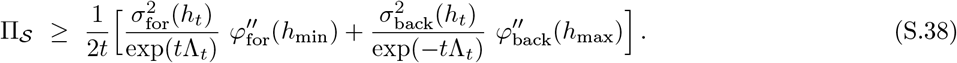

We notice that this version of the bound does not depend on the average values of the fitness landscape, unlike eq. (S.37).

Let us mention that if no information is known on the support of the fitness landscape, *h*_min_ can still be taken equal to 0, in both eqs. (S.37) and (S.38), because fitness landscapes are positive functions. Indeed, combining the fluctuation relations on *K* (eq. (1)) and *s* (eq. (2)) provides another expression for the fitness landscape [2]

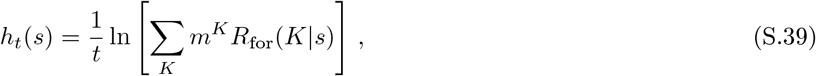

in terms of the conditional forward probability *R*_for_(*K|s*) of the number of divisions. Since *m ≥* 1 and *K ≥* 0, this relation implies that the fitness landscape is a positive quantity. In this case, eq. (S.37) (resp. eq. (S.38)) gives a non-trivial bound based on the first two moments (resp. second moment) of the forward fitness landscape distribution.

Note also that in the case where the support of the fitness landscape for the trait 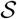 is unknown, but where the support of the number of divisions is known, then one obtains *h*_min_ ≥ *K*_min_ ln *m/t* and *h*_max_ ≤ *K*_max_ ln *m/t* from eq. (S.39).

### EXPERIMENTAL FITNESS LANDSCAPES

We tested our different inequalities on real data, extracted from [21], for the following traits: the number of divisions 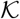, the size 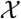 and the age 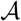, which are easily accessible and for which we predicted theoretical fitness landscapes [8].

We show the shapes of the fitness landscapes for 3 particular experiments on fig. 4. Each line corresponds to an experiment, the first column shows size fitness landscapes and the second one age fitness landscapes.

**Figure 4:**
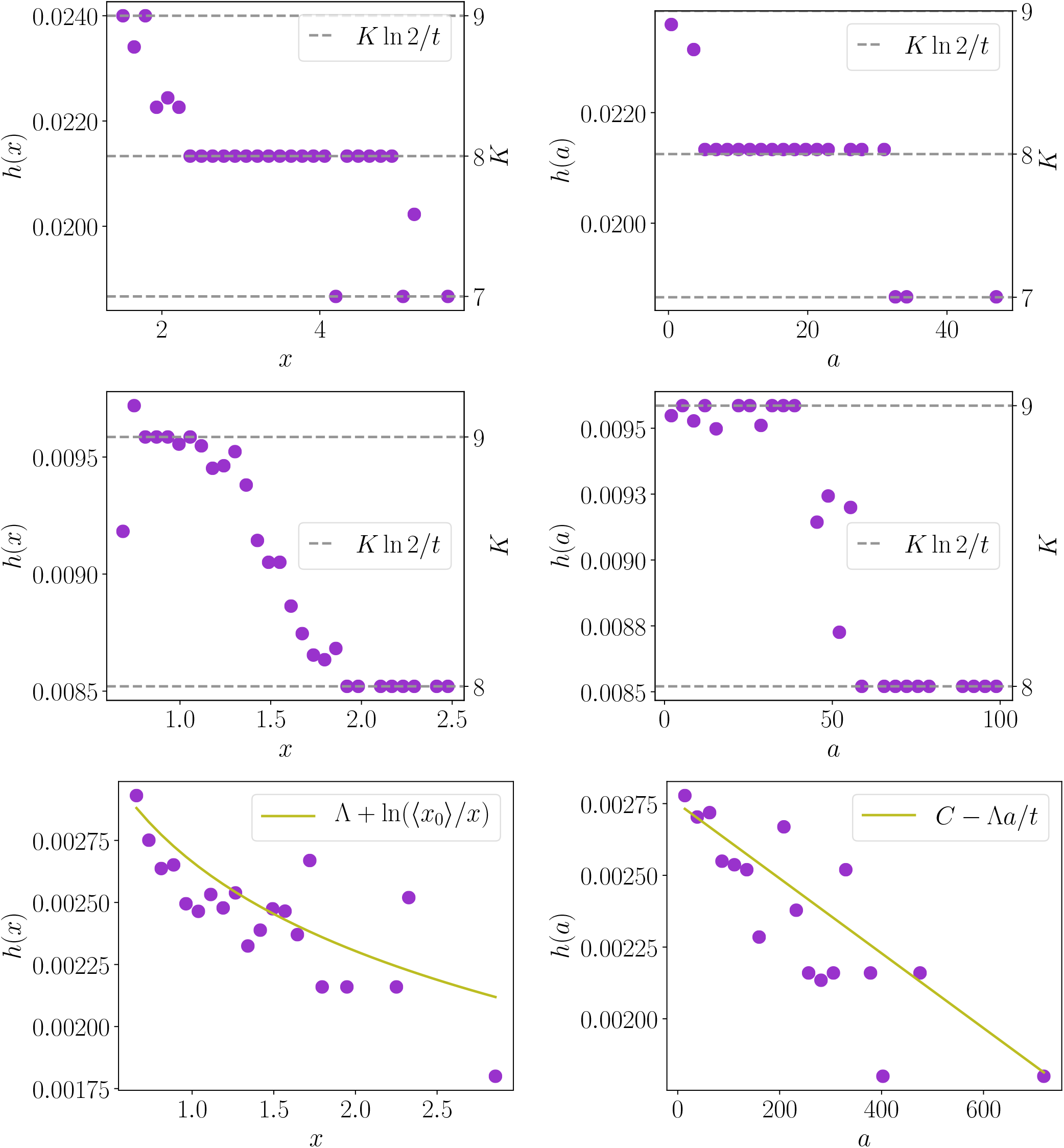
Experimental fitness landscapes computed with data from [21]. Each line corresponds to a different experiment, the first column shows fitness landscapes for size and the second column for age. For the first two lines, the grey horizontal dashed lines correspond to the plateaus predicted by the theory when *K* is fully determined by the value *s* of the trait. Plateau values are determined solely by the number of divisions undergone by the cell, and we see that for the same experiment, plateaus in size and age occur for the same numbers of divisions *K*. In the last line, the plateaus are blurred and the scatter plots show some agreement with the theoretical smooth curves computed for size, in the case where there is no variability in individual growth rate nor volume partition at division, and for age, in the case of a steady state [8]. The constant *C* was adjusted to fit the scatter plot, whereas no fitting parameter was needed for the size fitness landscape.

It is straight-forward to see that in the case where the number of divisions *K* is completely determined by the value *s* of the trait (so that *R*_for_(*K|s*) = *δ*(*K − K*(*s*)) in eq. (S.39)), then *h*_*t*_(*s*) = *h*_*t*_(*K*(*s*)) = *K*(*s*) ln 2*/t* [2, 8]. Thus the fitness landscape is an ensemble of plateaus corresponding to the values of *K* featured in the population at time *t*. Cells that are on the same plateaus have undergone the same number of divisions, even though having a different value *s*. Those predicted plateaus are actually observed for several experiments, as for the two first lines of fig. 4. The dots between the plateaus correspond to sizes/ages that have been reached by cells with different numbers of divisions. We see that the agreement between the predicted plateau values and the experimental data is very good, suggesting a strong control of the population.

As lineages de-synchronize because of the cumulative effect of various noises, we predicted that plateaus blur and are replaced by a smooth curve, whose equation we obtained under certain hypotheses for age and size in [8]. We predicted a logarithmic dependency in *x* for *h*_*t*_(*x*) and a linear dependency in *a* for *h*_*t*_(*a*), which are in agreement with the general shapes of the experimental fitness landscapes displayed on the third line.

